# Mutational landscape and *in silico* structure models of SARS-CoV-2 Spike Receptor Binding Domain reveal key molecular determinants for virus-host interaction

**DOI:** 10.1101/2020.05.02.071811

**Authors:** Shijulal Nelson-Sathi, PK Umasankar, E Sreekumar, R Radhakrishnan Nair, Iype Joseph, Sai Ravi Chandra Nori, Jamiema Sara Philip, Roshny Prasad, KV Navyasree, Shikha Ramesh, Heera Pillai, Sanu Ghosh, TR Santosh Kumar, M. Radhakrishna Pillai

## Abstract

Protein-protein interactions between virus and host are crucial for infection. SARS-CoV-2, the causative agent of COVID-19 pandemic is an RNA virus prone to mutations. Formation of a stable binding interface between the Spike (S) protein <u>R</u>eceptor <u>B</u>inding <u>D</u>omain (RBD) of SARS-CoV-2 and <u>A</u>ngiotensin-<u>C</u>onverting <u>E</u>nzyme 2 (ACE2) of host actuates viral entry. Yet, how this binding interface evolves as virus acquires mutations during pandemic remains elusive. Here, using a high fidelity bioinformatics pipeline, we analysed 31,403 SARS-CoV-2 genomes across the globe, and identified 444 non-synonymous mutations that cause 49 distinct amino acid substitutions in the RBD. Molecular phylogenetic analysis suggested independent emergence of these RBD mutants during pandemic. *In silico* structure modelling of interfaces induced by mutations on residues which directly engage ACE2 or lie in the near vicinity revealed molecular rearrangements and binding energies unique to each RBD mutant. Comparative structure analysis using binding interface from mouse that prevents SARS-CoV-2 entry uncovered minimal molecular determinants in RBD necessary for the formation of stable interface. We identified that interfacial interaction involving amino acid residues N487 and G496 on either ends of the binding scaffold are indispensable to anchor RBD and are well conserved in all SARS-like corona viruses. All other interactions appear to be required to locally remodel binding interface with varying affinities and thus may decide extent of viral transmission and disease outcome. Together, our findings propose the modalities and variations in RBD-ACE2 interface formation exploited by SARS-CoV-2 for endurance.

**Importance:** COVID-19, so far the worst hit pandemic to mankind, started in January 2020 and is still prevailing globally. Our study identified key molecular arrangements in RBD-ACE2 interface that help virus to tolerate mutations and prevail. In addition, RBD mutations identified in this study can serve as a molecular directory for experimental biologists to perform functional validation experiments. The minimal molecular requirements for the formation of RBD-ACE2 interface predicted using *in silico* structure models may help precisely design neutralizing antibodies, vaccines and therapeutics. Our study also proposes the significance of understanding evolution of protein interfaces during pandemic.

## Introduction

For SARS-like corona viruses, successful virus-host interaction initiates when RBD of virus S-protein binds to peptidase domain of host ACE2. This creates a favorable protein-protein interface for viral entry (Hoffmann et al., 2020, Ou et al., 2020). Being the first point of host contact, RBD has been a favorite target for vaccines, neutralizing antibodies and therapeutics against COVID-19 (Du et al., 2009). Recent crystal/ cryo-EM structures (Wrapp et al., 2020, Wang et al., 2020, Lan et al., 2020) and molecular models (Wang et al., 2020) of RBD-ACE2 complexes from SARS-CoV-2 and related SARS-CoV provided initial clues regarding molecular architecture of the interface. RBD comprises of 223 amino acid long peptide in the S1-region of S-protein (**Table 1**) (Walls et al., 2020). However, ACE2 binding information is confined to a variable loop region within RBD called <u>R</u>eceptor <u>b</u>inding <u>m</u>otif (RBM). These structures elucidated key interfacial interactions responsible for enhanced binding affinity of SARS-CoV-2 to ACE2 than SARS-CoV (Wang et al., 2020). It also suggested that few amino acid changes in RBM can remodel the interface resulting in altered binding affinities and viral transmission. However, all these studies were based on parental SARS-CoV-2 Wuhan strain and several questions remain unanswered. What are the mutations acquired on RBD during COVID-19 and what are the interfacial molecular rearrangements induced by these mutations? Can we gain valuable insights regarding RBD-ACE2 interface formation by analyzing these mutations? To address these questions we investigated the mutational landscape of SARS-CoV-2 RBD.

**Table 1:**
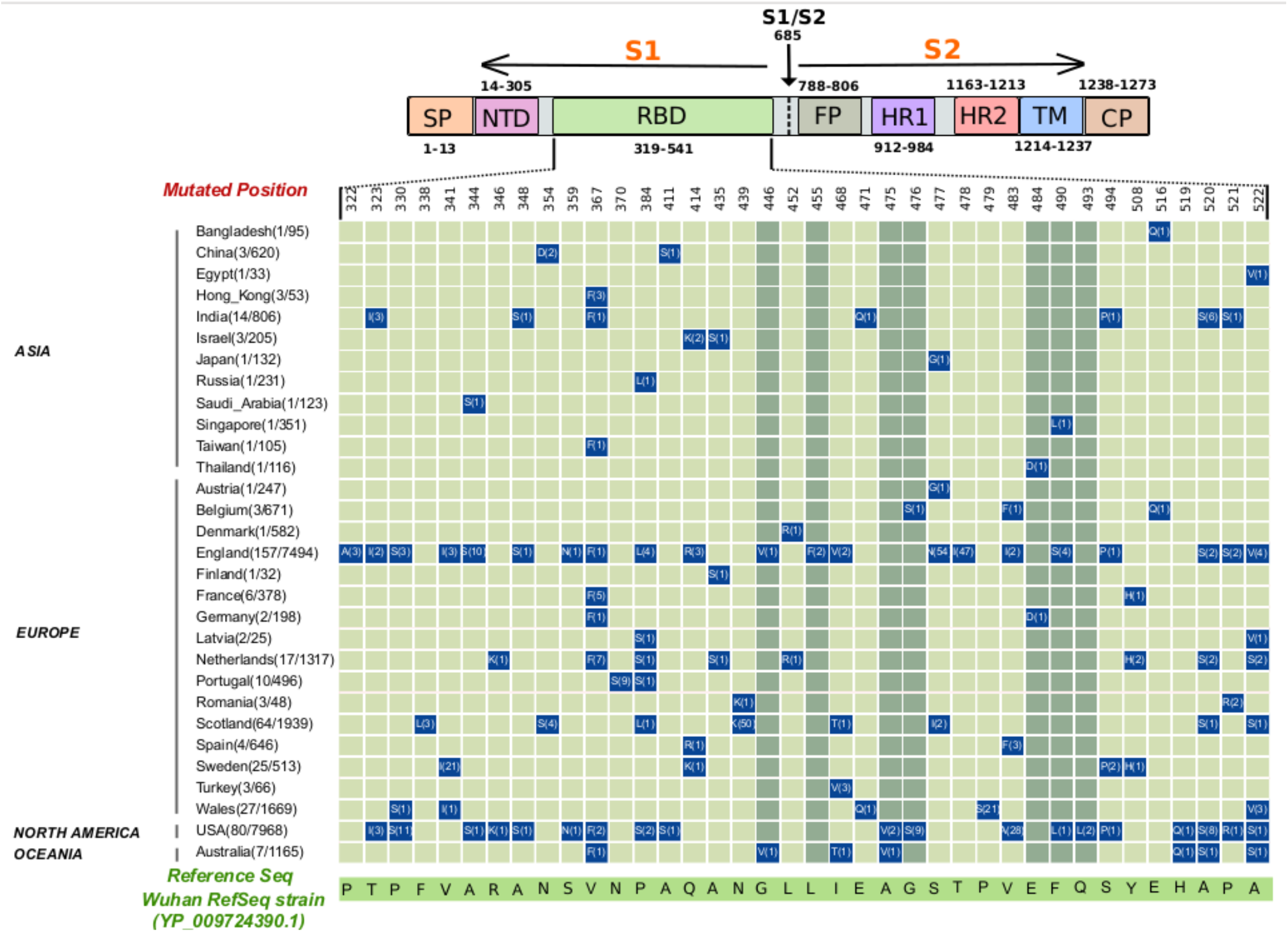
Matrix representing amino acid substitutions present in RBD domain of SARS-CoV-2 S protein of 31,403 genomes. Name of countries and the number of mutants *vs* genomes sampled are given on the Y-axis and the relevant amino acid residues (single letter code) in the reference strain are given on the X-axis. Mutated amino acid residues and their frequency of occurrence are provided in matrix cells. Light green colored matrix cells represent non-interface mutations and dark green color matrix cells represent interface-mutations in the RBD domain of spike protein. Mutations which are present at least in two independent genomes at the same position are represented in the matrix along with their positions.

## Methods

### Mutational analysis

A total of 55,485 spike proteins of SARS-CoV-2 were directly downloaded on 29^th^ June 2020 from the GISAID database. We removed the partial sequences, sequences greater than 1% unidentified ‘X’ amino acids and sequences from low quality genomes. Further, 31,403 spike protein sequences along with Wuhan reference spike protein (YP_009724390.1) were aligned using Mafft (*maxiterate* 1,000 and global pair-*ginsi*) (Katoh et al. 2002). The alignments were visualized in Jalview (Waterhouse AM et al., 2009) and the amino acid substitutions in each position were extracted using custom python script. We ignored the substitutions that were present in only one genome and unidentified amino acid X. The mutations that are present in at least two independent genomes in a particular position were further considered. These two criteria were used to avoid mutations due to sequencing errors. The mutated amino acids were further tabulated and plotted as a matrix using R script.

### Phylogeny reconstruction

For the Maximum-likelihood phylogeny reconstruction, we have used the SARS-CoV-2 genomes containing RBD mutations, and 10 genomes were sampled as representatives for each known subtype with Wuhan RefSeq strain as root. Sequences were aligned using Mafft *(maxiterate* 1,000 and global pair-*ginsi*), and phylogeny was reconstructed using IQ-Tree (Nguyen et al., 2015). The best evolutionary model (GTR+F+I+G4) was picked using the ModelFinder program (Kalyaanamoorthy et al., 2017).

### Structural analysis

The structural analysis of the mutated spike glycoprotein of SARS-CoV-2 RBD domain was done to assess the impact of interface amino acid residue mutations on binding affinity towards the human ACE2 (hACE2) receptor. The crystal structure of the SARS-CoV-2 RBD-hACE2 receptor complex was downloaded from Protein Data Bank (PDB ID:6LZG) and the mutagenesis analysis was performed using Pymol (DeLano WL et al., 2002). Homology modelling of Mouse ACE2 (mACE2) structure was performed in Swiss-Model (Schwede et al., 2003) using SARS-CoV-2 RBD-hACE2 as template. The YASARA server (Krieger et al., 2002) was used for the energy minimization of analysed structures. The Z-dock webserver (Pierce BG et al., 2014) was used for docking the mACE-2 and the spike protein RBD of SARS-CoV-2. The binding affinity of the wild, mutated and docked structures was calculated using PRODIGY web server (Xue, Li C., et al., 2016). The hydrogen bond and salt bridge interactions were calculated using Protein Interaction Calculator (Tina KG et al., 2007) and the Vander Waals interactions were calculated using Ligplot (Wallace AC et al., 1995). All the visualizations were done using Pymol (DeLano WL et al., 2002).

## Results and Discussions

### Non-synonymous RBD mutational profile

To capture changes that affect viral tropism during pandemic, we searched for non-synonymous mutations in RBD sequences from SARS-CoV-2 genomes. Using unbiased and stringent filtering criteria, we analyzed 31,403 genomes deposited in GISAID till 29th June, 2020. Altogether, 444 non-synonymous mutations in RBD were identified that belong to viral genomes from 30 countries. These mutations were found to substitute 49 amino acid residues in which 23 residues lie within RBM (**Table 1**). These residues include those that directly engage ACE2 (G446, L455, A475, G476, E484, F490 and Q493) and those that are in the near binding vicinity (**Figure 1A and 1B**). Hot spot mutations were also identified that caused recurrent substitutions of amino acid residues in the same position (N354, P384, Q414, I468, S477, V483, F490, A520, P521 and A522). Each RBD mutation was found to be unique to the genome; a combination of mutations was never observed in our analysis. Overall, RBD mutations accounted for ~9% of the total non-synonymous mutations in S-protein.

**Figure 1:**
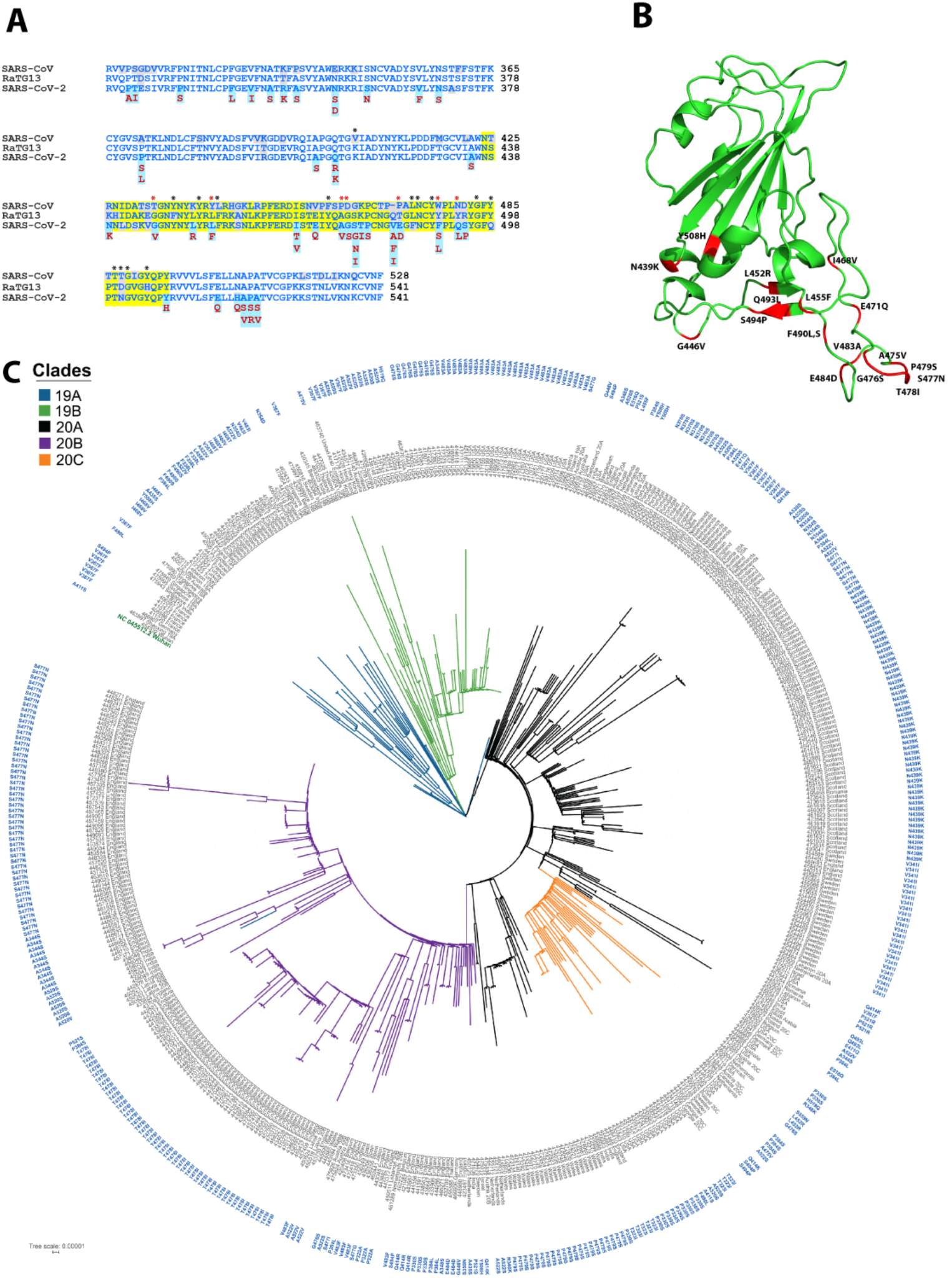
**A.** Conservation of Receptor Binding Domain (RBD) of SARS-CoV-2 with its close relatives, SARS-CoV and Bat RaTG13. The blue colored region shows RBD and the yellow highlighted region within RBD is the Receptor Binding Motif (RBM). The mutated residues are highlighted in light blue and substitutions are marked below. Nonconserved residues are in grey color. **B.** Maximum Likelihood Phylogenetic tree of 494 SARS-CoV-2 isolates containing RBD mutations. The outer circle represents the RBD mutations corresponding to each taxon marked in blue. Branches are coloured based on the known SARS-CoV-2 subtypes. The phylogeny is rooted with Wuhan RefSeq strain and highlighted in green.

### Evolutionary pattern of RBD variants

To see the evolutionary trend in RBD mutations, we compared RBDs from SARS-CoV-2, the related SARS-CoV and the bat coronavirus RaTG13, a suspected precursor of SARS-CoV-2. SARS-CoV-2 RBD is 73.4 % identical to SARS-CoV and 90.1 % identical to RaTG13 (**Figure 1A**). We identified several RBD mutations on residues that are unique to SARS-CoV-2 (N439K, V483A/F/I, E484D, F490S/L, Q493L and S494P) or are conserved in all three viruses (**Figure 1A**). In addition, we observed micro evolutionary reversion mutations in SARS-CoV-2 that interchange residues to that in SARS-CoV or RaTG13 (R346K, N354D, N439K, L452R, E471V and S477G) (**Figure 1A**). Interestingly, most of the unique and reversion mutations were located in the RBM region and thus may have implications in viral tropism. We performed phylogenomic analysis to understand the evolutionary pattern of RBD variants during pandemic and observed an unbiased clustering of RBD variants among distinct SARS-CoV-2 subtypes likely indicating independent emergence of these mutants (**Figure 1C**).

### Structural implications of RBD mutations

RBD is divided into a structured core region comprising five antiparallel β-sheets and a variable random coil region, RBM that directly binds to ACE2 (**Figure 1B**). On the contrary, binding information on ACE2 is located across long α-helices. Structurally, RBM scaffold resembles a concave arch that makes three contact points with ACE2 α-helix; Cluster-I, II and III. Cluster-I and Cluster-III are on two ends and Cluster-II is towards the middle of the interface (**Figure 2A**). We analysed the effect of observed RBM mutations on the molecular interactions at RBD-ACE2 binding interface. It has been shown that differences in ACE2 residues render mouse resistant to infection from SARS-like coronaviruses (Zhao et al., 2020). Hence, to gain insights into relevant interactions that can create stable interface in mutants, we also included RBD-mouse ACE2 interface in our analysis (**Figure 2B**). Structure models were created for all mutants based on the information from three recently reported crystal/ cryo-EM structures of SARS-CoV-2 RBD-ACE2 bound complex (**Figure 2A**). Comparative analysis of structures showed key differences in all three binding clusters of SARS-CoV-2 RBD wild type and mutant interfaces with human or mouse ACE2 (**Figure 2C, 2D and Table S1**).

**Figure 2:**
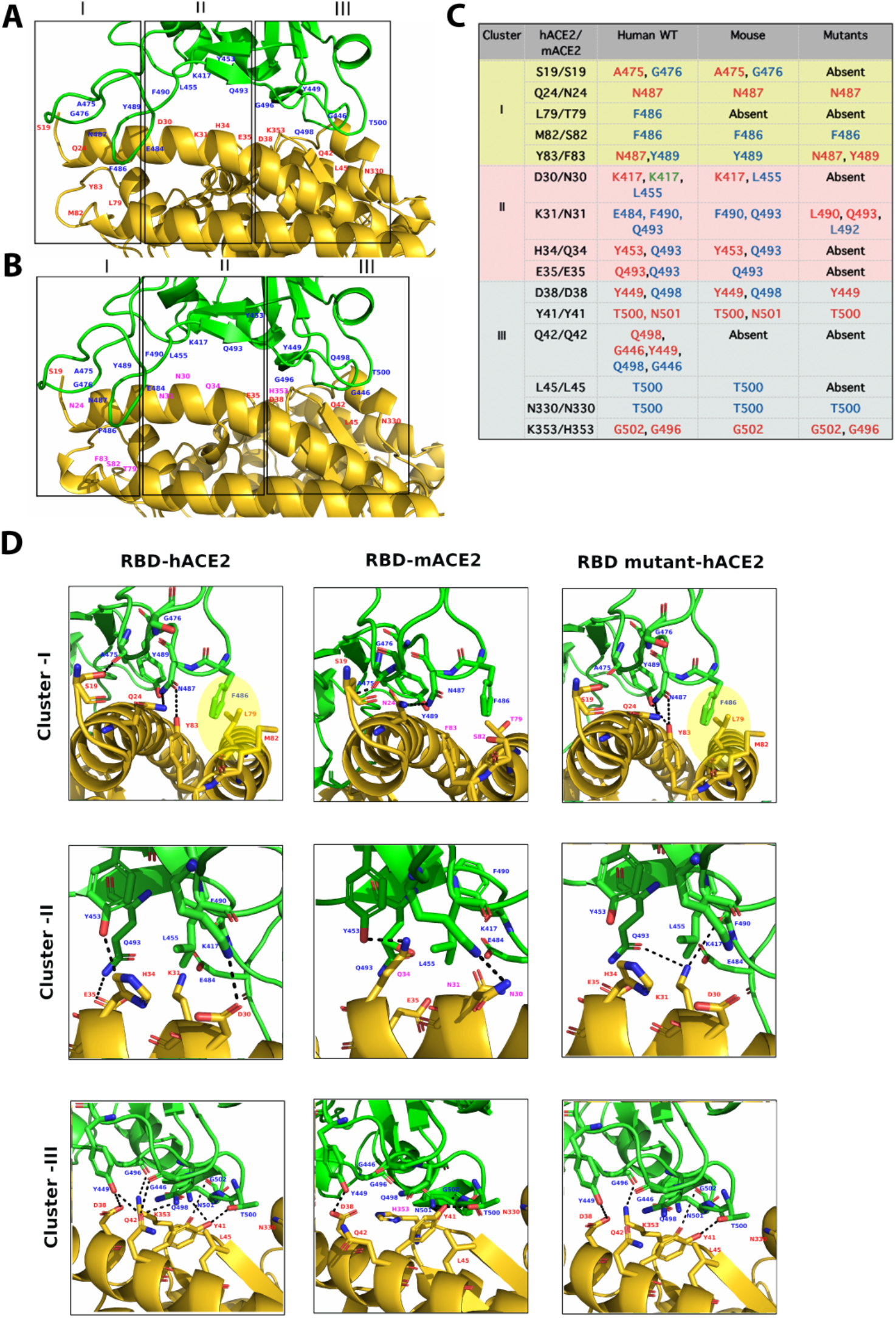
Molecular rearrangements in RBD-ACE2 interface. **A**. SARS-CoV-2 RBD-human ACE2 interface showing key residues (in red and blue) and clusters (boxed region) of interaction. RBD interface residue mutations identified are highlighted in yellow. **B**. SARS-CoV-2 RBD-mouse ACE2 docked interface. Mutated ACE2 residues in mouse are marked in purple **C**. List of cluster specific molecular interactions of hACE2, mACE2, mutated RBD-ACE2 complexes. Hydrogen bonds are marked in red, vander Waal’s in blue and salt bridges in green. **D-F.** Structural visualization of key interactions listed in **C**. SARS-CoV-2 RBD is represented in green and ACE2 in gold. The hydrogen bond interactions between ACE2 and RBD are shown as dotted lines.

In cluster-I, F486 of SARS-CoV-2 RBM is found buried into the hydrophobic pocket made of human ACE2 residues L79, M82 and Y83. A mutation of F>L in SARS-CoV disrupts this pocket, thus weakens the binding affinity suggesting importance of this interaction (Wan et al., 2020). In addition, N487 of SARS-CoV-2 RBM forms hydrogen bonds with Q24 and Y83 of human ACE2. The hydrophobic pocket and N487-Y83 interactions were completely abolished in mouse interface due to natural ACE2 substitutions in L79T, M82S and Y83F. But, these interactions were retained in all RBM mutants suggesting their importance in the pandemic. Nevertheless, interactions of A475/ G476 of SARS-CoV-2 RBD with S19 of ACE2 in cluster-I which were present in human and mouse were disrupted in mutants. In addition, SARS-CoV-2 genomes containing A475V and G476S replacements were identified in our analysis suggesting these mutations can be well tolerated. An additional hydrogen bond between Y489-Y83 was seen in mutants lacking A475/G476-S19 interaction. This could possibly be a compensatory mechanism to stabilize cluster-I interactions (**Figure 2C, 2D and Table S1**).

Cluster-II is predominated by polar and charged residue interactions including hydrogen bonds and salt bridges. A salt bridge formed between K417 (residue unique to SARS-CoV-2 and not present in SARS-CoV) and D30 of ACE2 has been reported to significantly enhance receptor binding in SARS-CoV-2 (Wang et al., 2020). Similarly, E484-K31, Q493-E35 bonds are also present in human to increase binding efficiency. These cluster-II interfacial interactions are abolished in mouse owing to ACE2 substitutions further underscoring their importance. Surprisingly, majority of RBM mutants fail to form these interactions including K417-D30 salt bridge. Nevertheless, compensatory hydrogen bond between Q493/L490-K31 was observed in these mutants (**Figure 2C, 2D and Table S1**).

A bunch of interactions in cluster-III involving G446/Y449/Q498 of RBM and Q42 of ACE2 were present in human but abolished in mouse and RBM mutants. However, additional interactions to compensate for these were not seen. A hydrogen bond formed between G496 of RBM and K353 of human ACE2 appeared significant as this was completely abolished in mouse, owing to K353H substitution, but retained in all RBM mutants. In addition, other interactions; G502-K353, Y449-D38 and T500-Y41 in the same cluster were maintained in human, mouse and mutants likely suggesting their supportive role (**Figure 2C, 2D and Table S1**).

The varying interface arrangements in mutants were consistent with the binding affinity differences (ΔΔG). Compared to wild type, ΔΔG values of mutants ranged within ~ + 1 kcal/mol, with the lowest value close to that of SARS-CoV (**Figure S1**). Sine RBM is a variable loop, mutations on any residue could impact spatial arrangements of backbone leading to altered binding affinities. Consistently, we did not find a considerable difference in binding energies between mutations on residues that are involved in ACE2 interaction or are in the near vicinity.

**Figure S1:**
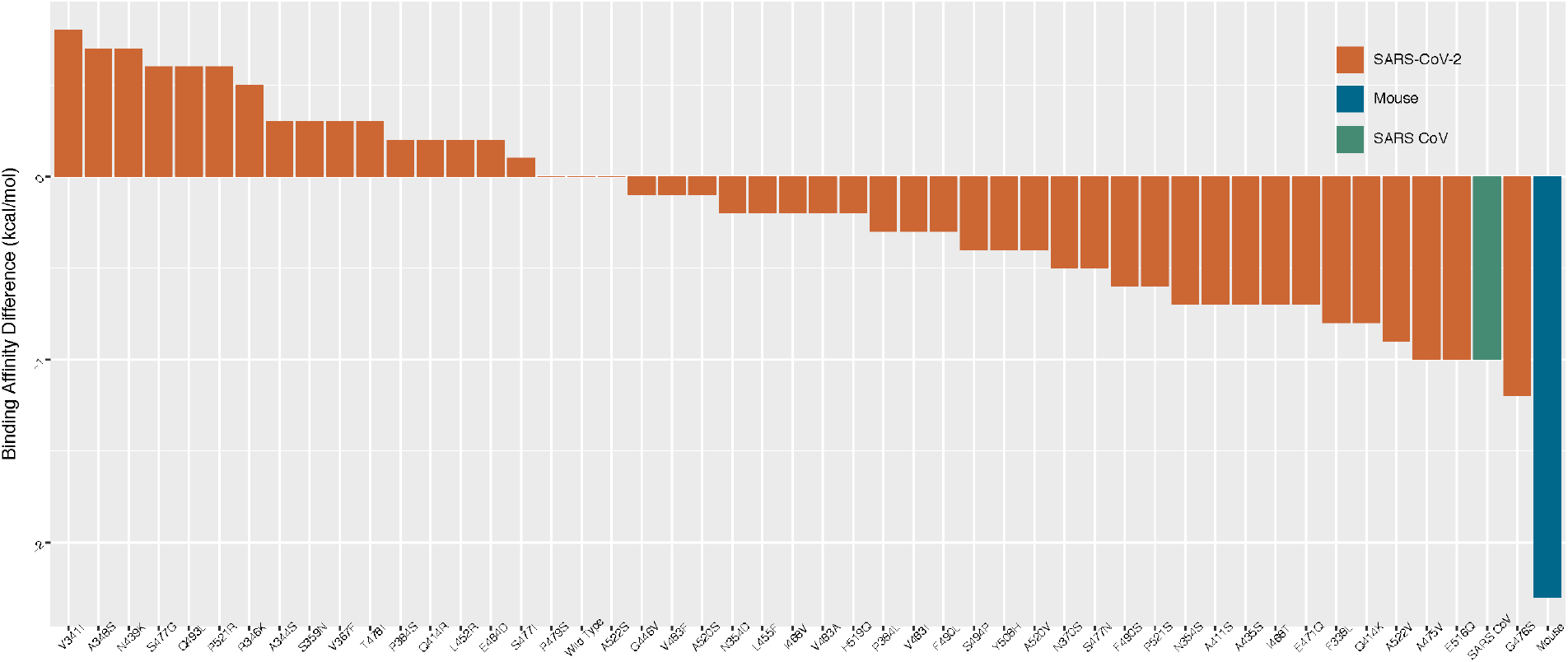
Bar graph representing variations in binding affinity differences among all RBD mutants. Values of Binding affinity difference (ΔΔG) is represented in kcal/mol on Y-axis and is calculated based on the net binding energy (ΔG) of the interface of parental SARS-CoV-2 Wuhan strain RBD with human ACE2. Bars corresponding to ΔΔG of SARS-CoV-2 mutants, SARS-CoV and mouse interfaces are highlighted in orange, green and blue colors respectively.

In conclusion, we could pinpoint two interfacial interactions that remain unaffected in all mutants analysed. These are interactions mediated through RBD residues N487 and G496 and are located in cluster-I and cluster-III respectively. Based on their spatial arrangement, these residues appear critical in directly anchoring the RBM loop onto ACE2. This may help initiate interface formation that favours viral entry. Both N487 and G496 are highly conserved in all SARS-like corona viruses further reinforcing this notion. The significant remodelling in cluster-II interactions indicates they are dispensable for anchoring but might be important for stabilizing the interface. Our findings are in accordance with the recent crystal structures (Wrapp et al., 2020, Wang et al., 2020, Lan et al., 2020) and molecular models (Wang et al., 2020) and provide additional mechanistic insights into formation of RBD-ACE2 interface.

Since RBD-ACE2 interface is a direct determinant of viral infectivity, along with other factors, varying interface architecture and binding affinities in SARS-CoV-2 RBM mutants may account for global variations in COVID-19 transmission and outcome. SARS-CoV-2 S-protein is highly immunogenic, so recombinant vaccines and neutralizing antibodies that target the whole S-protein or RBD are currently being considered in clinics (Zhang et al., 2020). Our investigations reveal key molecular determinants and their modalities for RBD-ACE2 interaction. This information might be used to design vaccines, synthetic nanobodies or small molecules that could specifically target RBM anchoring residues or their binding pockets.

## Acknowledgements

The authors wish to acknowledge John B Johnson, Mahendran KR and Sara Jones for critical comments. The work was supported by the Department of Biotechnology, Government of India.

